# Pulsatile basal gene expression as a fitness determinant in bacteria

**DOI:** 10.1101/2024.09.30.615870

**Authors:** Kirti Jain, Robert Hauschild, Olga O Bochkareva, Roderich Roemhild, Gasper Tkačik, Calin C Guet

## Abstract

Active regulation of gene expression, orchestrated by complex interactions of activators and repressors at promoters, controls the fate of organisms. In contrast, basal expression at uninduced promoters is considered to be a dynamically inert mode of non-functional “promoter leakiness”, merely a byproduct of transcriptional regulation. Here, we investigate the basal expression mode of the *mar* operon, the main regulator of intrinsic multiple antibiotic resistance in *Escherichia coli*, and link its dynamic properties to the non-canonical, yet highly conserved start codon of *marR* across *Enterobacteriaceae*. Real-time single-cell measurements across tens of generations, reveal that basal expression consists of rare stochastic gene expression pulses, which maximize variability in wildtype and, surprisingly, transiently accelerate cellular elongation rates. Competition experiments show that basal expression confers fitness advantages to wildtype across several transitions between exponential and stationary growth by shortening lag times. The dynamically rich basal expression of the *mar* operon, has likely been evolutionarily maintained for its role in growth homeostasis of *Enterobacteria* within the gut environment, thereby allowing other ancillary gene regulatory roles to evolve, e.g. control of costly-to-induce multi-drug efflux pumps. Understanding the complex selection forces governing genetic systems involved in intrinsic multi-drug resistance is crucial for effective public health measures.

---

Basal gene expression, also known as promoter leakiness, is a characteristic of bacterial promoters that occurs in their OFF state due to the presence of a repressor or the absence of an activator. Unlike induced or constitutive expression in the ON state, basal expression is generally not thought of as a functional mode of gene expression, but rather as a lack thereof. While selection can tune various aspects of gene induction, it is unclear how it could act, if at all, on the basal expression mode. Given that promoter leakiness can be detrimental (1–3), it could be under negative selection. However, we wondered whether there are any alternative basal expression modes that could have regulatory functions in their own right and thus be positively selected for. Recent studies uncovered the existence of a much more dynamic, pulsatile basal expression mode for several bacterial genes. Such a basal mode can generate phenotypic diversity in a clonal population and has thus been rationalized as a possible bet hedging mechanism (4–7). Essential to this explanation are two premises. The first is “frequency matching”: bet hedging conveys a long-term fitness benefit when it generates phenotypes in proportion to the frequencies of the environments for which these phenotypes are advantageous (8). The second is the existence of a growth rate “cost” for a pulse: some (small) fraction of cells undergoing a pulse pay this cost upfront, in order to survive, or be more competitive, if a rare external stress should occur in that moment. While the bet hedging explanation is attractive, the implied growth rate costs and benefits, as well as fitness effects more broadly, are rarely measured (4). This motivates a fundamental question: Are the two premises of bet hedging met or should one seek alternative explanations for the evolutionary maintenance of a pulsatile basal expression mode?

Here we turn to the *marRAB* operon, initially discovered as the genetic determinant of *m*ultiple *a*ntibiotic *r*esistance, and a paradigmatic example of a highly complex bacterial regulatory circuit (9, 10). The repressor MarR and the activator MarA form a negative and a positive autoregulatory loop, respectively, and this unique topology of two interlocked loops jointly controls the *mar* function (11, 12). While MarR is a local regulator of *marRAB* operon, MarA is a global regulator at the heart of one of the largest *E. coli* regulons, encompassing over 30 genes, involved in multi-drug efflux, pH regulation, outer membrane permeability, biofilm formation, and virulence (13–18). Modelling and experimental studies suggest that the *mar* interlocked regulatory loops could lead to pulsatile basal mode, resulting in phenotypic heterogeneity of expression that could support bet hedging (11, 19). Nevertheless, the characterization of the basal expression mode for the *mar* operon and its functional and fitness implications beyond bet hedging and antibiotic stress remains unexplored.

## Conservation of GTG start codon in *marR* across *Enterobacteriaceae*

As MarR controls the repressed state of the *mar* operon and therefore its basal expression, the unusual presence of a weak GTG start codon in *marR* piqued our interest (20). Non-ATG start codons, i.e., GTG and TTG, initiate ∼8% genes in *Gammaproteobacteria* (21), reducing the translation efficiency of these genes so that significantly lower expression is achieved than if genes used ATG. To determine whether the GTG start of *marR* is a historical contingency or the outcome of selection, we constructed the *marR* phylogenetic tree and determined GTG prevalence across *Gammaproteobacteria*. Among 889 representative genomes, *marR* homologs were found in ∼300 species, distributed across 20 distinct bacterial families (**Fig. 1A**). Interestingly, we observed that *marR* belongs to the *marRAB* operon only in *Enterobacteriaceae*. Within all other bacterial families, *marR*-type transcription factors form operons with *emrAB*-type efflux pump genes **(Fig. 1A, S1)**.

**Fig. 1.**
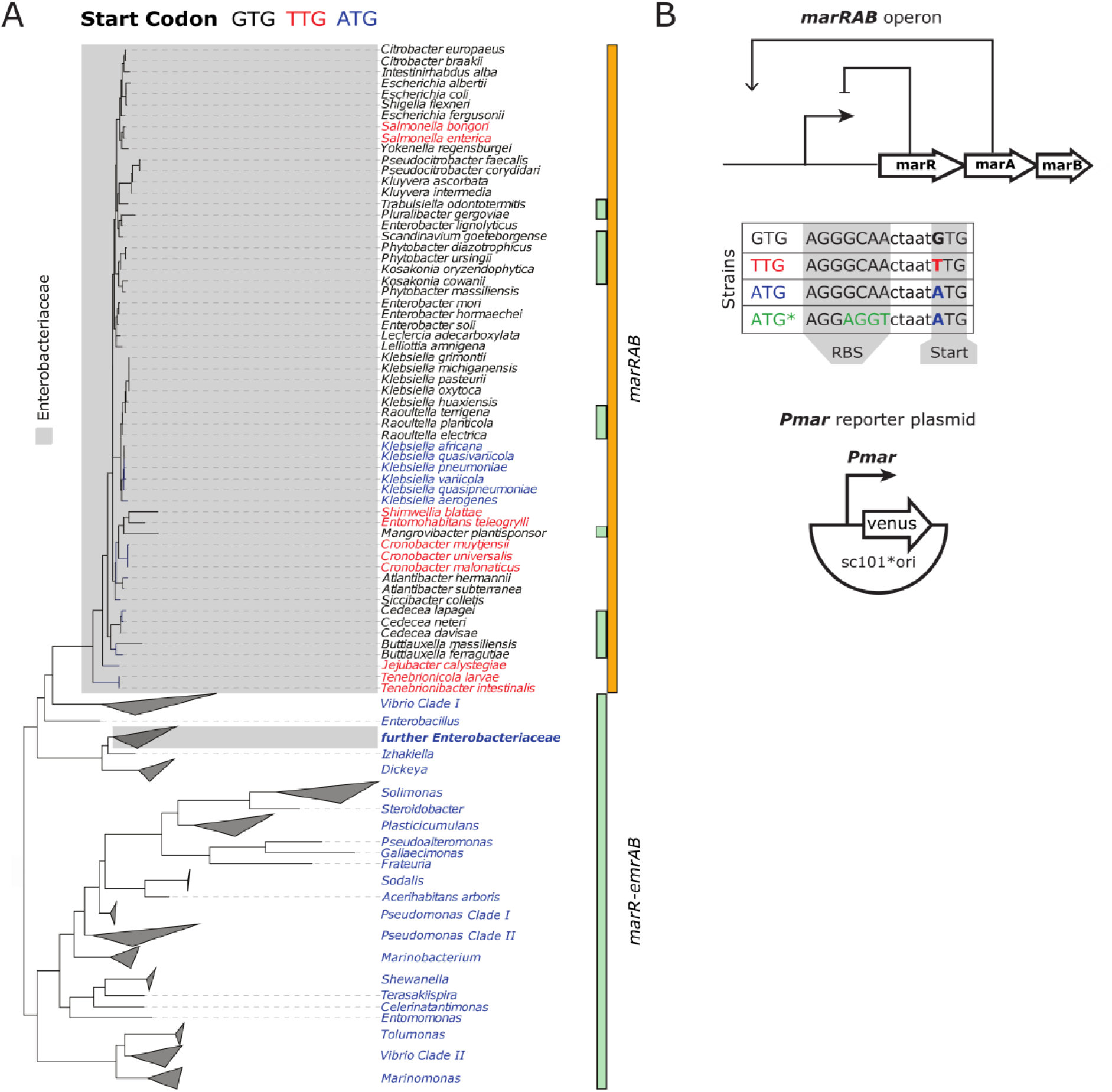
Distribution of *marR* start codons across *Enterobacteriaceae*. **(A)** Maximum likelihood phylogenetic tree of *marR* in *Gammaproteobacteria*. The tree was constructed using other members of *marR* family (full tree in **Fig. S1**). The *marRAB* operon is only present in *Enterobacteriaceae*, while in other bacterial families, a *marR*-type transcription factor was found in conjunction with the genes forming an *emrAB*-type efflux pump. **(B)** Schematic representation of *marRAB* operon (top), strains constructed (middle), and reporter plasmid (bottom).

The phylogenetic tree corroborates an evolutionary scenario in which *marRAB* operon evolved only once and was vertically inherited **(Fig. 1A)**. Its formation in the ancestor of *Enterobacteria* coincides with a change of the *marR* start codon from the canonical ATG to the noncanonical GTG. While the *marR-emrAB* family has a strong ATG start codon, *marR* in *marRAB* operons uses the weaker GTG variant, with very few exceptions (*Cronobacter*, *Jejubacter, Pluralibacter, Salmonella, Shimwellia*, *Tenebrionocola*), where the putatively even weaker TTG is used. Furthermore, a switch from GTG to ATG occurred only in one *Klebsiella* clade. Few *Enterobacteria* harbor both, *marRAB* and *marR-emrAB* operons, (*Cedecea*, *Kosakonia*, *Phytobacter*, *Raoultella*), providing evidence for horizontal gene transfer of *marR-emrAB* into some *Enterobacteriaceae* (**Fig. 1A**). In addition to the start codon, the ribosome binding site (RBS) is also a determinant of translational efficiency. With a single exception, the RBS of *marR* in *marRAB* is fully conserved among *Enterobacteria* (**Fig. S2**). Taken together, our phylogenetic analysis strongly suggests that the prevalent utilization of weak *marR* start codons across *marRAB* operons, in conjunction with a particular RBS variant, is selectively favored for a yet uncharacterized, but likely general, physiological role.

### Pulsatile basal expression of *mar* operon

To ask how the conserved GTG start codon affects *marRAB* function and fitness, we constructed scarless *marR* mutants with alternative start codons (ATG and TTG) in *Escherichia coli*. In addition, in the ATG* mutant we combined the ATG start codon with a stronger RBS **(Fig. 1B)**.

To investigate the basal *mar* expression mode in single cells, we measured the fluorescence output of a *Pmar*-*venus* promoter fusion using time-lapse microscopy in a microfluidic device (22, 23). We simultaneously monitored a chromosomal constitutive *P_R_*-*mCherry* as a control. Average background-corrected *Pmar* expression depended significantly on the choice of start codon **(Fig. 2A)**. TTG yielded 3-4 fold higher *Pmar* expression than the wildtype (GTG), consistent with the expectation that TTG leads to weaker repressor translation. In contrast, the strong, canonical ATG start codon reduced *Pmar* basal expression below the wildtype (GTG) levels. The ATG* mutant that combines the canonical ATG start codon with a strong RBS, abolished most of *Pmar* expression and accessed a nearly complete OFF promoter state **(Fig. 2B)**.

**Fig. 2.**
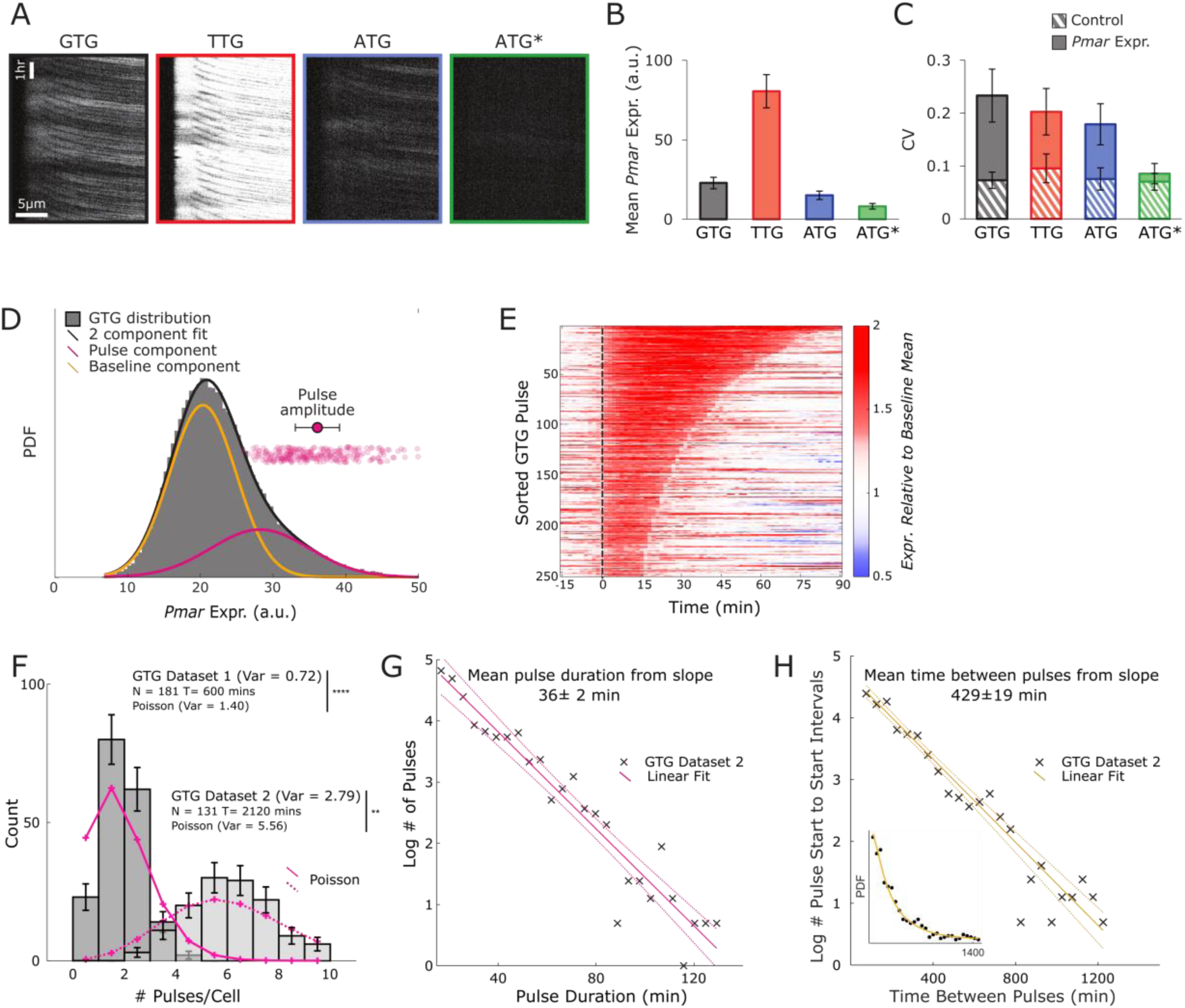
Characterization of *Pmar-venus* basal expression dynamics. **(A)** Kymographs of wildtype (GTG) and mutants (TTG, ATG, and ATG*) showing *Pmar-venus* expression for a representative mother cell imaged in a microfluidic channel over 10 hours. **(B)** Mean *Pmar-venus* expression in wildtype and mutants (Mean and SD over cells, p<10^-4^, rank-sum test). **(C)** Coefficient of variation (CV) of *Pmar-venus* expression in wildtype and mutants compared against CV of control (constitutive reporter *P_R_-mCherry* expression). *Pmar-venus* expression CV in wildtype is significantly different from mutants (mean and SD over cells, p<10^-4^, rank-sum test). (*Pmar-venus* expression CV is also significantly different from the corresponding *P_R_-mCherry* control for each strain, p<10^-4^, rank-sum test). **(D)** Distribution of *Pmar-venus* expression is modeled as a sum of two Gaussian distributions, i.e., baseline and pulse component, for wildtype. Magenta circles represent the amplitudes of individual extracted pulses and magenta circle with an error bar is the mean amplitude ± SD over individual pulses. **(E)** Pulses for wildtype sorted by duration and pulse start is aligned to 0 min on the x-axis. **(F)** Distribution of the number of pulses per cell in wildtype in Dataset 1 (181 cells, 600 mins) and Dataset 2 (131 cells, 2120 mins) showing significant deviation from Poisson. Error bars represent √(Count). **(G)** Log histogram of pulse durations. Mean pulse duration (in caption) was derived from the inverse of the slope of the linear fit. **(H)** Log histogram of time intervals between two consecutive pulses. Inset shows the corresponding PDF.

We next characterized the overall variability of the basal expression mode by computing the coefficient of variation (CV) of *Pmar-venus* fluorescence across 10 hours of observation for each of the ∼180 independent mother cells per genotype **(Fig. 2C)**. The wildtype (GTG) showed maximum expression variability, followed by TTG (despite having higher mean expression than that of the wildtype), and then ATG. These three strains have at least 2-fold higher variability than constitutive controls. For ATG*, the CV was only slightly elevated relative to the control **(Fig. 2C, S3)**.

The observed high *Pmar* variability in single cells traces its origin to gene expression pulses: transient, stochastic, high-amplitude activations of transcription plainly visible in all strains (**Movies M1-M5**). To extract and statistically characterize these pulses **(Fig. S4),** we first decomposed the observed *Pmar-venus* fluorescence distributions into a Gaussian mixture. The frequent lower-amplitude component corresponded to baseline fluctuating *Pmar* expression levels, whereas the rarer component corresponded to sporadic high-amplitude pulse-like excursions **(Fig. 2D, S5)**. This motivated a 85-percentile threshold (see Methods) for extracting the pulses which could subsequently be aligned to their respective start times **(Fig. 2E, S6)** and quantified.

We report a similar frequency of pulsing in the wildtype (GTG), TTG and ATG strains, of one pulse per approximately 7-8 hours. For ATG*, pulses appear to be much less frequent, but our detection may be biased by their low signal-to-noise ratio **(Table 1)**. The pulse duration distribution was exponential for the wildtype (GTG), TTG, and ATG strains for which it could be reliably estimated, with similar average duration of ∼33-37 minutes per pulse **(Table 1, Fig. 2G, S7)**. The key difference between the strains lay in the overall *mar* expression that affects the baseline as well as pulse amplitudes **(Table 1, Fig. S8)**. This expression changed several-fold depending on the MarR start codon, implying that the MarR translation efficiency can tune *Pmar* expression by determining the promoter activity level outside and during the pulse. When expressed as fold-change increase over their respective baseline Gaussian components – which we refer to as pulse “signal-to-noise” ratio (SNR) – differences between strains were smaller but significant: pulse amplitudes ranged from ∼1.3 – 1.7, with the maximal SNR reached in the wildtype (GTG) **(Table 1, Fig. S8A).** This difference at the level of pulse characteristics is responsible for the maximal CV in *mar* expression reported for the wildtype (GTG). Finally, after z-scoring and accounting for the individual durations of the pulses, we find that pulses nearly collapse onto a universal shape, indicating that most of the variability across genotypes is accounted for by the statistics we extracted **(Fig. S8B).**

**Table 1.** Characterization of pulse expression, amplitude, duration, and frequency in wildtype (GTG), TTG, ATG, and ATG* strains.

|  | GTG | TTG | ATG | ATG* |
| --- | --- | --- | --- | --- |
| Baseline mean expression (a.u.) | 20.35±4.56 | 71.57±13.53 | 13.49±3.02 | 7.36±1.04 |
| Pulse mean SNR (amplitude / baseline mean) | 1.69 ±0.01 | 1.59 ± 0.01 | 1.55 ± 0.01 | 1.33 ± 0.02 |
| Baseline mean relative to Wildtype | 1.00 | 3.50 | 0.66 | 0.36 |
| Pulse amplitude relative to Wildtype | 1.00 | 3.39 | 0.61 | 0.30 |
| Pulse mean duration (min) | 35±20 | 37±23 | 33±16 | 25±11 |
| Pulse frequency (per h) | 0.14 | 0.12 | 0.15 | 0.04 |

Can pulsing be modeled as a stationary stochastic point process? If that were the case, pulse numbers should be Poisson-distributed over individual cells of the same genotype. We report strong and highly significant deviations from this expectation for the wildtype (GTG), TTG, and ATG strains but not for ATG* and controls **(Fig. 2F, S9),** even though inter-pulse intervals are exponentially distributed for all strains **(Fig. 2H)**. Specifically, the observed pulse count distributions are under-dispersed compared to Poisson, suggesting a more regular pulsing, possibly due to finite pulse duration or pulse-pulse correlations. Despite these quantitative deviations, individual pulses could be interpreted in the bet-hedging framework as stochastic switches into an alternative (high *marA*) phenotype once every ∼14 – 16 generations and lasting for about one generation.

We next assessed the cost of *mar* basal mode expression. TTG cells had a significantly lower long-term elongation rates than wildtype (GTG) cells, whereas the elongation rates of ATG and ATG* were marginally higher than for the wildtype (GTG) **(Fig. 3A)**. This corresponds to the ordering of mean *mar* expression levels across strains **(Fig. 2B)** and is consistent with the expectation that higher overall *mar* activity is costly. A detailed analysis, however, revealed a surprising finding. We compared the single-cell long-term elongation rates to the instantaneous elongation rates during different phases of the pulse. We expected the elongation rates to slow down around a pulse and subsequently return to the long-term average. In contrast, for all strains but ATG* we observed significantly increased elongation rates in the time window 0-20 minutes after we identify the pulse start **(Fig. 3B, C)**. The growth advantage could be caused by differences in pulse amplitudes, where different sets of *mar* regulon targets are engaged by different levels of MarA. This selective targeting is plausible, since genes in the *mar* regulon are known to respond continuously and with different sensitivities to MarA levels (24). Taken together, larger baseline *mar* expression has a cost, while a rare transient pulse confers an advantage. Therefore, selection may have to navigate this tradeoff in an environment-dependent way.

**Fig. 3.**
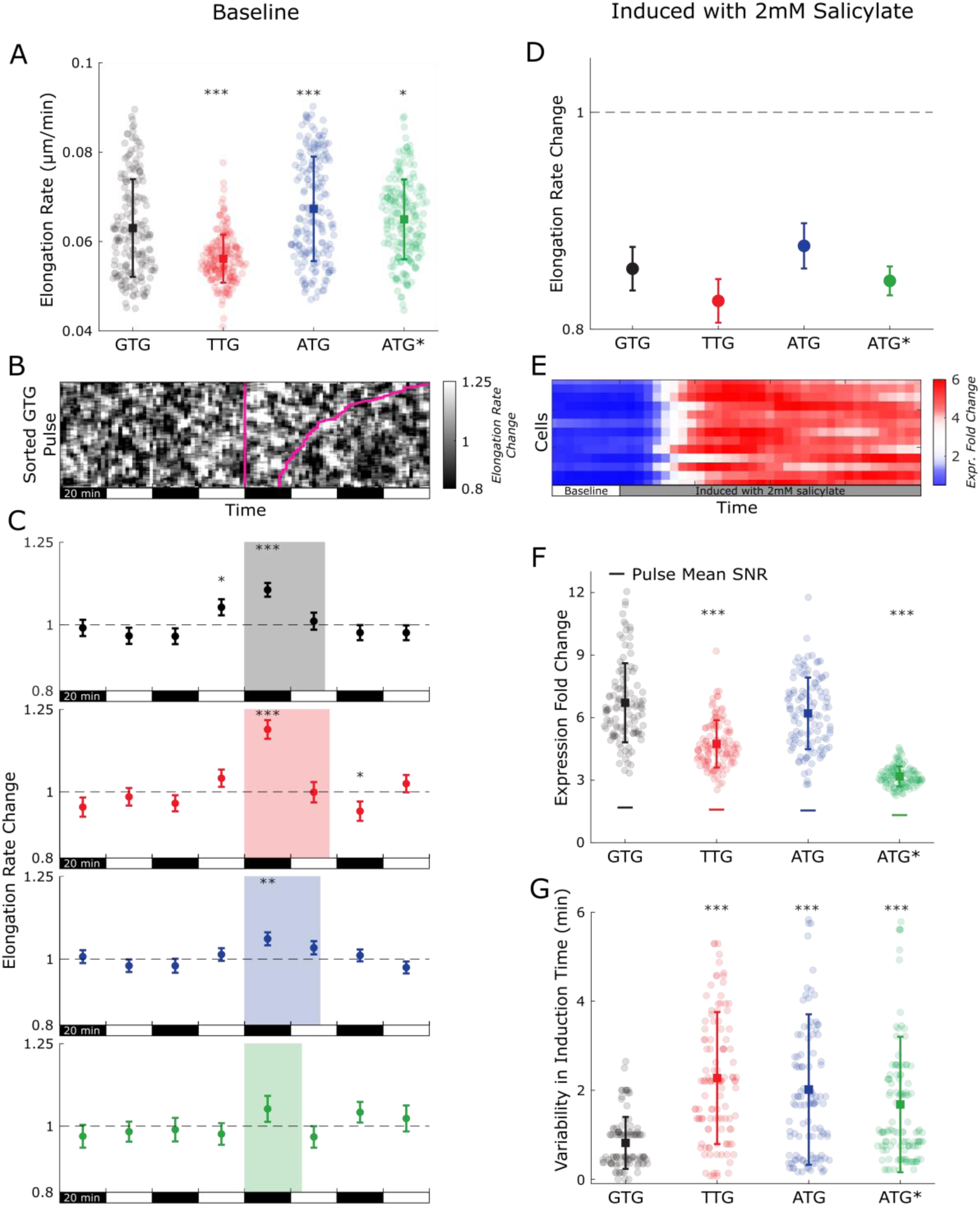
Elongation rates and *Pmar-venus* expression quantification in baseline (A to C) and induced state (D-G). **(A)** Single-cell elongation rates in the baseline state (individual cell data (circles) with mean (square) and SD). Wildtype elongation rate is significantly different from TTG & ATG (p<10^-4^, rank-sum test) and ATG* (p<0.05, rank-sum test). **(B)** Raw data showing instantaneous elongation rate normalized to the long-term individual cell average and aligned to pulse start (vertical magenta line). Pulse duration is denoted in magenta as well. **(C)** Mean and SE of elongation rates (normalized to the long-term average elongation rate of the respective cell) in 20 min windows (80 mins before and after the pulse start) for wildtype and mutants. The shaded regions illustrate the start of the pulse and the mean pulse duration in each strain. Stars indicate significant deviation from the long-term elongation rate (t-test). **(D)** Mean and SE of elongation rate change upon induction with 2mM salicylate relative to their respective baselines shows similarly-sized fold change relative to pre-induction baseline and statistically significant reduction across strains (p<10^-4^, t-test). **(E)** Representative raw data (*Pmar-venus* expression) showing the transition from baseline to 2mM salicylate-induced state. **(F)** Individual cell data (circles), mean (square), and SD of fold change of *Pmar-venus* expression upon induction with 2mM salicylate, horizontal line shows a comparison with pulse mean SNR in the baseline state. Wildtype expression fold change upon induction is significantly different from TTG & ATG* (p<10^-4^, rank-sum test) and ATG (<0.05, rank-sum test). **(G)** Individual cell data (circles), mean (square), and SD of variability in induction time quantified by the time taken by expression to reach the inflection point for each cell. Wildtype is significantly different from all three mutants (p<10^-4^, rank-sum test).

To verify that low and transient (as opposed to high and persistent) *mar* expression during the pulse is necessary for an elongation rate advantage, we exposed ∼110 cells per strain in our microfluidic device to 2mM salicylate to induce *Pmar* expression **(Fig. 3E).** Induction caused a prolonged increase in *marRAB* expression, with the largest, ∼6-7 fold induction in the wildtype (GTG) and ATG, followed by TTG (4-5 fold) and ATG* (∼3 fold) **(Fig. 3F)**. In terms of absolute expression, these levels were substantially (2x to 4x, depending on the strain) above the pulse amplitudes in the basal mode **(Table 1).** Induction brought about a concomitant ∼10% decrease in the elongation rate in all strains **(Fig. 3D)**, consistent with previous reports and our expectation that prolonged and strong *marRAB* expression is detrimental likely because it engages additional, more costly-to-express *mar* regulon targets (24).

The induction experiment revealed another significant difference between the wildtype (GTG) strain and the strain with the canonical start codon (ATG), which emerged when we analyzed the inflection times of individual cell induction curves **(Fig. 3G)**. The wildtype (GTG), which exhibits maximal expression variability in the basal mode, surprisingly had the least variable induction curves and thus the most synchronized response to induction. In comparison, the mutants show a two-fold reduction in synchronicity. Overall this suggests that the unique interlocked regulatory circuit with short-lived MarA results in precise induction to sudden stress (25), indicating that response speed and synchrony may be at a premium for the enterobacterial ecological niche. The observed difference between the wildtype (GTG) and the canonical start codon (ATG) strain motivated us to focus next on the fitness effects of *mar* in environments where *mar* expression mode transitions and timing could be relevant.

### Fitness advantage for the wildtype basal expression mode across growth cycles

The surprising observation that basal mode pulse brings about a transient elongation rate advantage suggests a role of *mar* expression in physiology and growth homeostasis, which is in line with the subtle influence of *mar* in the transition from exponential to stationary phase and back (26, 27). We therefore decided to conduct pairwise competition experiments to assess the performance of our strains across the entire growth cycle. We competed the wildtype (GTG) vs ATG or vs ATG* strains, and vs GTG itself (as a control), across four serial growth cycles over four days (together > 40 generations) in LB media without any external inducers. The key question was how ATG and ATG*, the two strains with more efficient MarR repressor translation and thus lower baseline *mar* expression, compare against the wildtype (GTG) when the cells are forced to undergo repetitive transitions between exponential, stationary, and lag phases.

Starting from a 1:1 ratio, we saw the wildtype (GTG) increase to a ratio of 2:1 over the course of 53 generations in competition with the ATG strain; the same 2:1 ratio was reached in 35 generations in competition with ATG*. The effective selection coefficients were –0.013 and –0.020 per generation for ATG and ATG*, respectively, relative to wildtype (GTG) **(Fig. 4A** and **S10).** The fitness advantage of the wildtype (GTG) in the absence of external inducers implies a functional role of the conserved weaker GTG start codon for cell physiology during serial growth cycles, likely via its effects on the pulsatile basal mode of *mar* expression. To understand how the wildtype (GTG), which is at a disadvantage during exponential growth, outcompeted the two mutant strains, we measured how quickly various strains recovered from the lag phase and transitioned to exponential growth. We report a delay of around 8-12 minutes for ATG and ATG* mutants vs the wildtype (GTG) in LB medium that matched conditions used in the competition experiment. The delay further increased by up to 25-50 minutes for nutrient poor M9 media **(Fig. 4B)**.

**Fig. 4.**
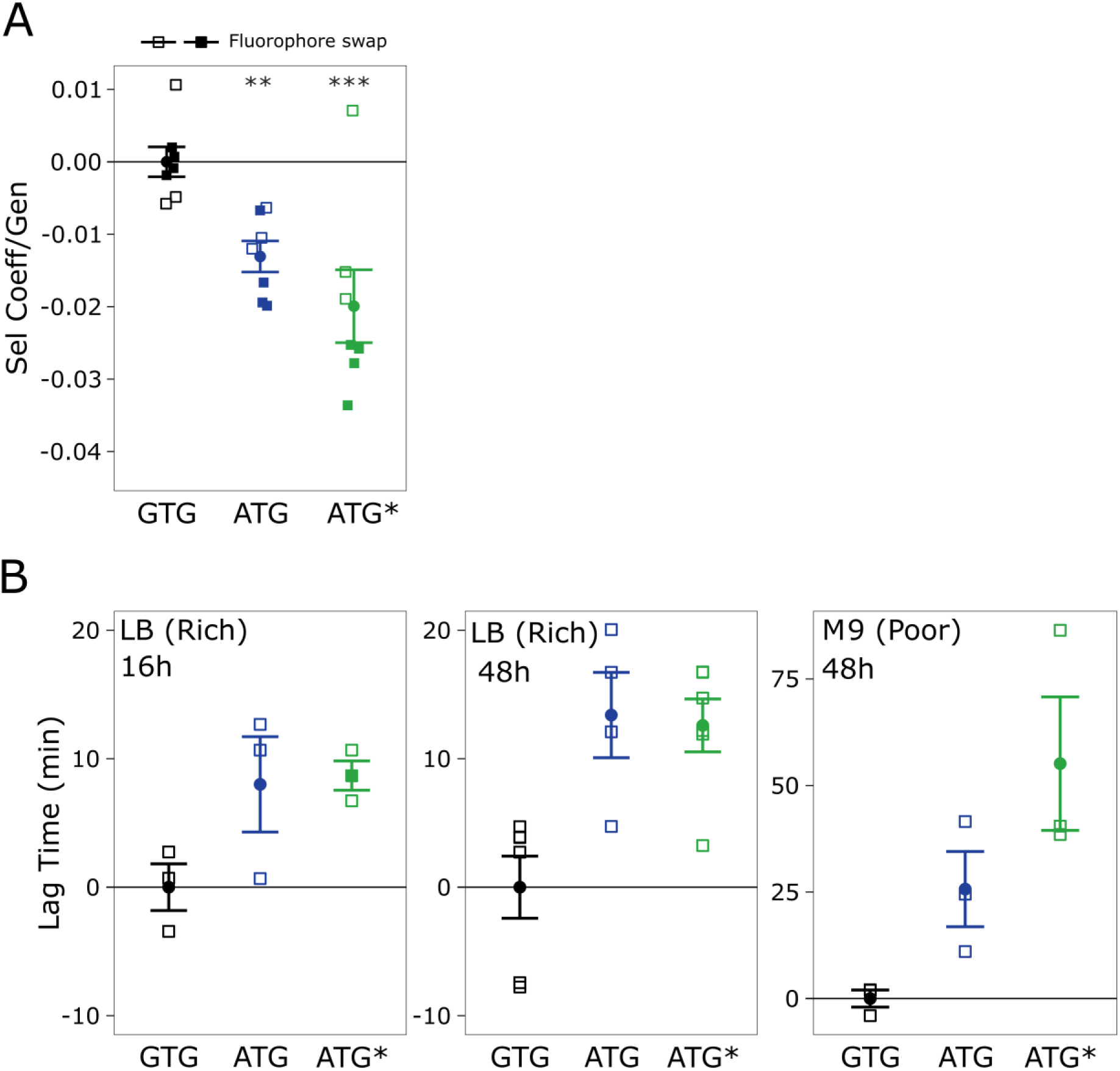
Competition and regrowth dynamics of wildtype (GTG) over ATG and ATG* mutants. **(A)** Selection coefficient (square) was quantified from the slope of the log ratios of competitor over wildtype strain per generation (circle represents mean and error bar represents SE over squares). ATG and ATG* lost the competition against the wildtype as determined by negative selection coefficients that were significantly different from the wildtype-wildtype control competition (s = –0.01, p<10^-2^ for ATG; s = –0.02, p<10^-4^ for ATG*, post hoc test and GLM) but not significantly different from each other (s = 0.007, post hoc test and GLM). mCherry (empty squares) and Venus fluorophores (filled squares) were used as markers to select for the respective strain background, and seven biological replicates across two independent experiments, including fluorophores swaps, were performed for each combination. **(B)** Time to regrow from stationary phase. Data points (square), mean (circle), and SE of lag time to initiate exponential growth when revived from 16 hours overnight LB (fresh rich media) culture, 48 hours LB culture (aged rich media), and 48 hours old M9 glycerol culture (aged poor media). Analysis of variance of lag time delays for the three experimental regrowth conditions showed a significant effect of strain (F = 9.4, p<10^-4^, ANOVA) and a significant effect of regrowth condition (F = 7.8, p<10^-2^, ANOVA). Sample size was three to five biological replicates per condition.

In summary, these results suggest that the wildtype (GTG) strain compensates for its slower exponential growth rate by shortening its lag time. It is instructive to consider a simple back-of-the-envelope calculation using realistic estimates from Fig. 3. If the long-term exponential growth rate of the wildtype (GTG) is 5% slower than that of the ATG mutant but it emerges from lag phase with a 24 min advantage, then the ATG strain would require 8 hours of uninterrupted growth to compensate for its delay and reach the same population size as the quicker-to-emerge but slower-to-grow wildtype strain. While *Enterobacteria* growth cycles are certainly more complex in the wild than in our lab setup, this simple estimate shows that the strategy of shortening the lag time could be competitive in practice.

## Discussion

The process of turning genes ON and OFF is fundamental to life, and regulatory networks that control it have been the object of intense study in developmental, evolutionary, molecular, and systems biology. Nevertheless, the question of whether the properties of the basal expression in the OFF state of promoters can be selected for was rarely, if at all, considered. We showed that the *mar* operon basal expression mode is highly dynamic, consisting of pulsatile gene expression at the single cell level, and we measured the fitness consequences of such dynamics. To this end, we used synonymous codon mutations for the start codon of *marR*, which we uncovered to be evolutionarily conserved and tightly coupled to the interlocked regulatory architecture of the *marRAB* operon.

We find that the weaker, non-canonical, GTG start codon acts as a regulatory knob ensuring the presence of sporadic transcription pulses that are much less pronounced or absent if *marR* and *marA* are translated with similar speed, as is the case in the various start codon mutants we measured. The GTG start codon endows the wildtype basal expression mode with several unique characteristics: highest expression variability due to highest “signal-to-noise” ratio of individual pulses, transient growth advantage during a pulse, and synchronized expression during transition to the induced state. Altogether, these quantitative expression characteristics lead to a robust fitness advantage for the wildtype that becomes apparent in environments in which cellular physiology switches between exponential, lag phase, and stationary growth. More broadly, this points to the involvement of *marRAB* operon in general growth homeostasis.

While the *marRAB* operon can be induced by various metabolic intermediates, no main physiological inducers have been characterized so far (28). However, the presence of nearly identical *mar* systems across most *Enterobacteria* and the remarkable conservation of the non-canonical GTG start codon of *marR* suggest that the cellular processes controlled by the *mar* operon confer adaptations that are most likely linked to *Enterobacteriaceae* physiology and their ecological niche (29). Essential to this ecological niche is that bacteria spend a significant fraction of their existence inside the guts of various animals. The quasi regular intervals of feeding that are characteristic of the gut environment point towards a selective pressure for optimizing cycles of lag, exponential, and stationary phases. Our results support the idea that the *mar* pulsatile basal expression mode could be an essential building block of this physiological adaptation to the intestinal lifestyle, allowing the basal expression mode to be evolutionarily maintained.

A supporting observation for this hypothesis is the quantitative match between the multiple timescales related to the *mar* operon in *Escherichia coli*: the typical time between two *mar* pulses, the timescale at which the shorter lag of wildtype would balance out its slower growth compared to *marR* start codon mutants, and the daily feeding cycles, which are all on the order of ∼10 hours. In the bet hedging framework, this could represent a quantitative case of frequency matching: yet instead of thinking of very rare stresses (such as antibiotic exposure), the system has evolved to match the frequency of environmental transitions typical for the ecological niche of *Enterobacteriaceae*. Constant selective pressure to maintain the *mar* basal expression mode for such physiological *raison d’etre* would enable the same system to be co-opted for other hypothesized functions in its induced mode: to generate diversity necessary for bet-hedging against much rarer stresses, or to induce costly stress-resistance genes when needed, and thus allow MarA to become a global transcriptional regulator. In addition to cyclic nutrient availability, the gut ecology is also characterized by host defense mechanisms and antimicrobials secreted by other microbiota. Thus, co-opting regulation of physiological adaptation to cyclic nutrient availability and antibiotic resistance mechanisms would be consistent with the global regulatory role of *mar* (30).

The naming of the *mar* operon – an acronym for *m*ultiple *a*ntibiotic *r*esistance coined over 40 years ago by George and Levy (9, 31) – was based on the conferred broad antibiotic resistance phenotype in the induced state (32). By now, it is clear that the *mar* operon has not evolved “for” antibiotic resistance, which is at best an ancillary function – albeit one of fundamental importance for public health. To control it and counteract the looming multi-drug resistance epidemic, our work demonstrates the acute need to understand the role of *marRAB* operon in bacterial physiology and growth homeostasis, in line with Seoane & Levy who argued early on for an alternative role of *mar* as a conveyor of ‘multiple adaptational response’ (10).

## Materials and Methods

### Computational Genomics

Genomes of *Gammaproteobacteria* were downloaded from RefSeq database (33) using the PanACoTA pipeline (34), module ‘download’ with filters: genome collections = ‘Reference’ or ‘Representative’; assembly level = ‘complete genomes’ or ‘chromosome’ or ‘scaffold’. Then we used the PanACoTA module ‘annotate’ to predict and functionally annotate CDS in the genomes.

*marR* genes were found using HMM for OG #1S2AX comprising *E.coli marR* (UniProt entry ID:P27245), from eggNOG5 database with e-value threshold 10^-35^ (35, 36). We additionally used HMMs for OG #1S26B and OG #1RPCJ which contain *slyA* (UniProt entry ID:P0A8W2) and *mprA* (UniProt entry ID:P0ACR9), correspondingly to include into the analysis more members of *marR* family in order to better resolve the gene tree **(Fig. S1)**.

To find ribosomal binding sites (RBSs) and correct putative errors in gene start prediction, *marR* upstream regions of 100 bp length were extracted and aligned taking into account the known RBS of *marR* in *E. coli* (20). Four downstream genes were used for the annotation of the *marR* genomic context.

The alignments of protein sequences and gene upstreams was made using Muscle v. 3.8.31 (37). The gene trees were constructed using IQtree v.1.6.12 with 1000 bootstrap runs (38). Visualization and annotation of the trees was done using Itol server (39). The logo of *PmarRAB* upstream was created using WebLogo 3 web-based application (40).

### Strains and Media

All experiments were performed using the derivate of *Escherichia coli* K-12 MG1655 strain, with incubations at 37°C and aeration.

Lysogeny Broth (LB) media was used in all experiments except when noted. For plates, 1.5% agar was added to LB. For microfluidics experiment, 0.01% Tween 20 was used to prevent attachment of bacterial cells to the PDMS device. Media and antibiotics were from Sigma, Sylgard for making PDMS was from Dow Chemicals. List of strains and primers used in this study are listed in **(Table ST1, ST2)**.

### Strain Construction

The DIRex method was used to generate scar-free point mutations for changing the start codon and RBS for *marR* (41). Briefly, the method uses a single λ Red recombineering step (pSIM5-Tet temperature sensitive) and a semi-stable AcatsacA intermediate. The desired changes were introduced through custom-made oligos with a homology to the target region. The AcatsacA cassette codes for three genes which help in selection and counter selection steps a) *cat* leading to chloramphenicol resistance (use 12.5 mg/L chloramphenicol to select for AcatsacA formation), b) *amilCP*, present as dual inverted copies, causing AcatsacA+ colonies to be of blue color helping in selection, and c) *sacB*, sucrose sensitivity gene (use 5% sucrose to select for self-excision), helps in counter selection generating scar-free mutants.

We use a constitutive chromosomal *P_R_-mCherry* reporter as control (22). *Pmar-venus* reporter is on a low copy plasmid (23).

### Microfluidics Set Up and Imaging

We used the same microfluidic chip as previously used in our lab (22). The length of the growth channel is 24µm and the width of these growth channels ranged from 1.2µm to 1.4µm and the height is approximately 1.1µm. To make mother machine devices from Epoxy replica, we used PDMS in a ratio of 10:1 (Sylgard and curing agent) and mixed and degassed in Thinky Machine (THINKY ARE-250) for 2 minutes each. Further degassing was done after pouring on the epoxy replica using a desiccator. Curing of PDMS was done overnight in an incubator at 80°C. Next, the PDMS device was peeled out carefully from the epoxy replica and holes were punched using an electro polished 18ga needle. The device was cleaned with scotch tape and the cover slip (24mm x 50mm, thickness 0.17mm+/-0.005) was cleaned with isopropanol. Device was then bonded using plasma bonding technique (Harrick PDC-002 plasma cleaner, medium power for 1 min, for both PDMS and cover slip) followed by gently placing on the cover slip. After bonding, it was kept on a hot plate (∼80°C) for one hour.

Before starting the experiment, the device was wetted with 0.01% Tween 20 for a couple of minutes followed by blowing out. This step also ensures that the bonding is leak-proof. Next, a pellet from exponential grown cells (overnight culture of the desired genotype in LB plus Tween 0.01% is grown and sub-cultured 1:1000 and grown for around four hours and centrifuged at 4000 X*g* for three minutes) were loaded using a pipette. After confirming the loading of cells by checking under the scope, media flow was connected with polyethylene tubing (BTPE –50Instech). Image acquisition settings were kept identical throughout all experiments (Exposure time for *mCherry*: 200ms and *venus*: 300ms) with an image interval of 90 secs. Images were acquired with an Olympus IX83 inverted fluorescence microscope, a 100X NA 1.45 objective, with a custom made autofocus, and a Hamamatsu Orca Flash4.0v2 camera (42).

### Image Analysis

#### Pre Segmentation

Images of the channel areas within the microfluidic chip were cropped and background and shading corrected (42).

#### Segmentation

Bacteria segmentation was carried out using Cellpose (43). A custom model was trained on a dataset of over 2000 hand-labeled cells selected from a diverse range of expression levels and morphologies.

#### Tracking

A customized Matlab script was employed for tracking, generating lineage trees, and conducting further analyses. In the initial stage, the microfluidic channels were automatically detected, and cells located outside the channels were eliminated. A heuristic method was applied to address missing cell detection and correct under-segmentation errors. Subsequently, each channel underwent individual tracking: The link cost function that establishes connections between cell detections at consecutive time points to form tracks, took into account the specific characteristics of the mother machine. This was achieved by assigning a higher link cost to reverse movement, instances where cells swapped positions within the channel, and a reduced overlap in cell segmentation compared to the segmentation at the previous time point. Cell divisions were identified when two cells overlapped with the same segmentation from the preceding frame. Empty channels were omitted for clarity.

#### Growth rate

We defined and quantified the growth rate and the promoter activity as done by Kim et al. (6, 44). The growth rate *g* is calculated as the logarithm of the ratio of the area of the cell immediately prior the cell division *A*_*d*_ and the area at the initial time point following the last cell division 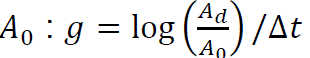.

As we did not observe any long term change in the average cell size, and we set 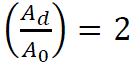.

According to this definition, it can be inferred that the growth rate remains constant between individual cell divisions.

#### Instantaneous elongation rate and cell division events

The instantaneous elongation rate R is calculated as 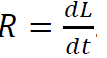. We derived the length *L* from a linear fit of the cell segmentation area, rather than an ellipse fit to the segmented cell, as this method was more robust against segmentation errors. To smooth out the growth rate, we applied a running average over a 15-minute window. Cell division events were identified by employing criteria that recognize when the segmented area of a cell approximately halves during division. Cell division events were excluded from the growth rate analysis.

#### Pulse detection and Analysis

The distribution of combined raw fluorescence intensities is accurately modeled as a sum of two distinct Gaussian distributions. This modeling provides a natural cutoff value, effectively differentiating baseline expression fluctuations from stochastic pulse-like expressions. Using this cutoff value, we perform a z-transform on each cell’s raw fluorescence intensities, utilizing the mean and standard deviation of the baseline expression. Pulses are identified by applying a threshold to the z-score **(Fig. S4)**. This method is employed for both fluorescence derived from the *Pmar-venus* promoter fusion and the chromosomal constitutive *P_R_-mCherry* expression. Additionally, we conducted a stochastic simulation replicating the characteristics of the constitutive expression. Analysis of the pulse length histogram in all three cases supports a cutoff for determining the duration of genuine gene expression pulses: Pulses shorter than 15 minutes are deemed random baseline fluctuations, whereas longer pulses, indicative of actual gene expression, are compiled for further examination. A linear regression on the inverse slope of the histogram of true pulses provides an average pulse length comparable to the directly calculated mean pulse duration. We also fitted Poisson distributions to the number of pulses per cell.

For visualization, pulses are aligned temporally, setting the start time of each pulse to t=0 and sorting them by duration. This approach allows for the calculation of the average intensity over time from *Pmar-venus* by normalizing each pulse’s intensity to its peak value. However, the transient pulse intensity is affected by two factors: the transient nature of each pulse and the distribution of pulse duration. To isolate the impact of varying pulse duration, we normalize the duration of each pulse to one. Consequently, the ensemble average of all duration-normalized pulses reveals the stereotypical shape of a pulse.

#### Inflection point

In order to quantify the synchronicity of induction, the inflection point of the fluorescence *I* was determined by finding the peak of 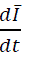 with 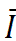 being the ten min moving average of *I*. The induction magnitude *M*, (= fold change of *I*) is given by the ratio of the average *I* at a fixed time interval before and after the inflection point:

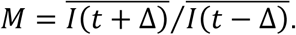

#### Statistical tests

We performed rank-sum tests for pairwise comparison of wildtype (GTG) with mutants for *Pmar-venus* and *P_R_-mCherry* expression. For elongation rate during pulse and elongation rate upon induction, we performed t-tests.

### Competition Experiment

We used two different marker methods: fluorophores (mCherry and Venus) **(Fig. 4A)** and resistance (chloramphenicol) **(Fig. S10)**. Each genotype was grown overnight in four replicates for each marker separately. Competitions were set up as head-to-head competitions of two genotypes, with dye-swap controls, where each marker was used for half of the replicates. The optical density (OD) was measured and pairwise cultures (wildtype (GTG): wildtype (GTG), wildtype (GTG):ATG, wildtype (GTG):ATG*) were mixed in a 1:1 ratio, diluted 1000x as to permit for ten generations of growth and incubated at 37°C for 24 hours. The 1000x-fold dilution was repeated for three consecutive days so that the genotypes were in direct competition for a total of 40 generations. In parallel, the cultures were plated on LB agar for CFU quantification of the genotypes. Plates were imaged using a custom-build fluorescence macroscope and fluorescent colonies were counted using ImageJ (42). When using the resistance markers, cells were plated on both LB agar and LB agar with chloramphenicol. Selection coefficients were calculated as the slope of the linear model fit to the log ratio of genotypes (using natural log) over generations of competitive growth. We corrected for fitness costs of the markers by subtracting the baseline selection coefficients of control competitions where the same genetic background was competed against itself with different markers. We then used a generalized linear model with genotype as fixed factor and experiment as random factor and a post-hoc test with user-defined contrasts to statistically test for significant selection between genotype pairs and estimate the selection coefficient. These analyses were done using the statistical software *R* and the packages *multcomp* and *nlme*.

### Population measurements of growth rate and lag phase

Growth rate and growth lag were measured by strongly diluting overnight cultures into 0.3 mL of fresh media in Honeycomb plates and measuring OD during exponential regrowth at 37°C with vigorous shaking every 4 min using the Bioscreen C plate reader (OY Growth Curves, Helsinki, Finland; Ref. FP-1100-C) system for at least 6 hours. Exponential growth rates were estimated as doublings per hour using the slope of the linear model fit to the plot of log-transformed OD over time in hours during the exponential growth phase. Accuracy of exponential growth was assess using R^2^ values of the fit, which was >0.94 for all growth curves (average 0.98). Lag phase was estimated as the time to restart exponential growth of OD, for which we used the time at which OD reached above the detection threshold of OD = 0.004. Time delay of mutant strains over the wild type strain was calculated by subtracting the average lag time of the wild type from the lag times of the mutants. To allow for estimation of lag phase growth curves were started with equal ODs. ODs of overnight cultures were normalized using OD immediately before inoculation into prepared Honeycomb plates. Without this correction, start growth time is dependent on starting cell density (45). We used ANOVA to test for significant differences in growth parameters between strains and three different growth conditions, where “LB, fresh” refers to regrowth in LB following dilution of a 16h overnight culture in LB, “LB aged” refers to regrowth in LB following dilution of a 48h culture in LB and “M9 glycerol aged” refers to regrowth in M9 minimal medium with 0.2% glycerol as the sole carbon source following dilution of a 48h culture in that medium.

## Author contributions

Conceptualization: KJ and CCG

Methodology: KJ (microfluidics, image analysis, competition and regrowth experiments), RH (analysis script), OOB (computational genomics), RR (competition and regrowth experiments)

Investigation: KJ (microfluidics, image analysis, competition and regrowth experiments), OOB (computational genomics), RR (competition and regrowth experiments)

Visualization: KJ, RH, OOB, RR, GT, CCG

Funding acquisition: CCG

Project administration: KJ and CCG

Supervision: GT and CCG

Writing – original draft: KJ

Writing – review & editing: KJ, GT, CCG, RR, OOB, RH

## Competing interests

Authors declare they have no competing interest.

## Classification

**Major** – *Biological Sciences* and **Minor** – *Systems Biology, Microbiology*

## Acknowledgements

KJ thanks B. Wu, I. Tomanek, K. Tomasek for detailed discussions on the manuscript, all other members from the Guet laboratory for helpful feedback, and R. Chait, & IOF IST Austria for helping with the microscope.

## Funding

KJ acknowledges IST fellowship IC1006FELL02, RH was supported in part by CZI grant DAF2020-225401 (10.37921/120055ratwvi) from the Chan Zuckerberg Initiative DAF, OOB acknowledges FWF Grant ESP253-B, RR acknowledges FWF Grant 10.55776/ESP219, CCG acknowledges FWF I5127-B.

## Data availability

All data to understand and assess the conclusions of this research are available in the main text and supplementary file. Raw data is at the IST repository.

## Notes

### Competing Interest Statement

The authors have declared no competing interest.

